# Scalable parameter estimation for genome-scale biochemical reaction networks

**DOI:** 10.1101/089086

**Authors:** Fabian Fröhlich, Barbara Kaltenbacher, Fabian J. Theis, Jan Hasenauer

**Affiliations:** Helmholtz Zentrum München - German Research Center for Environmental Health, Institute of Computational Biology, 85764 Neuherberg, Germany; Technische Universität München, Center for Mathematics, Chair of Mathematical Modeling of Biological Systems, 85748 Garching, Germany; University of Klagenfurt, Institute of Mathematics, A-9020 Klagenfurt, Austria

## Abstract

Mechanistic mathematical modeling of biochemical reaction networks using ordinary differential equation (ODE) models has improved our understanding of small-and medium-scale biological processes. While the same should in principle hold for large-and genome-scale processes, the computational methods for the analysis of ODE models which describe hundreds or thousands of biochemical species and reactions are missing so far. While individual simulations are feasible, the inference of the model parameters from experimental data is computationally too intensive. In this manuscript, we evaluate adjoint sensitivity analysis for parameter estimation in large scale biochemical reaction networks. We present the approach for time-discrete measurement and compare it to state-of-the-art methods used in systems and computational biology. Our comparison reveals a significantly improved computational efficiency and a superior scalability of adjoint sensitivity analysis. The computational complexity is effectively independent of the number of parameters, enabling the analysis of large-and genome-scale models. Our study of a comprehensive kinetic model of ErbB signaling shows that parameter estimation using adjoint sensitivity analysis requires a fraction of the computation time of established methods. The proposed method will facilitate mechanistic modeling of genome-scale cellular processes, as required in the age of omics.

**Author Summary:** In this manuscript, we introduce a scalable method for parameter estimation for genome-scale biochemical reaction networks. Mechanistic models for genome-scale biochemical reaction networks describe the behavior of thousands of chemical species using thousands of parameters. Standard methods for parameter estimation are usually computationally intractable at these scales. Adjoint sensitivity based approaches have been suggested to have superior scalability but any rigorous evaluation is lacking. We implement a toolbox for adjoint sensitivity analysis for biochemical reaction network which also supports the import of SBML models. We show by means of a set of benchmark models that adjoint sensitivity based approaches unequivocally outperform standard approaches for large-scale models and that the achieved speedup increases with respect to both the number of parameters and the number of chemical species in the model. This demonstrates the applicability of adjoint sensitivity based approaches to parameter estimation for genome-scale mechanistic model. The MATLAB toolbox implementing the developed methods is available from http://ICB-DCM.github.io/AMICI/.

## Introduction

In the life sciences, the abundance of experimental data is rapidly increasing due to the advent of novel measurement devices. Genome and transcriptome sequencing, proteomics and metabolomics provide large datasets [1] at a steadily decreasing cost. While these genome-scale datasets allow for a variety of novel insights [2, 3], a mechanistic understanding on the genome scale is limited by the scalability of currently available computational methods.

For small-and medium-scale biochemical reaction networks mechanistic modeling contributed greatly to the comprehension of biological systems [4]. Ordinary differential equation (ODE) models are nowadays widely used and a variety of software tools are available for model development, simulation and statistical inference [5, 6, 7]. Despite great advances during the last decade, mechanistic modeling of biological systems using ODEs is still limited to processes with a few dozens biochemical species and a few hundred parameters. For larger models rigorous parameter inference is intractable. Hence, new algorithms are required for massive and complex genomic datasets and the corresponding genome-scale models.

Mechanistic modeling of a genome-scale biochemical reaction network requires the formulation of a mathematical model and the inference of its parameters, e.g. reaction rates, from experimental data. The construction of genome-scale models is mostly based on prior knowledge collected in databases such as KEGG [8], REACTOME [9] and STRING [10]. Based on these databases a series of semi-automatic methods have been developed for the assembly of the reaction graph [11, 12, 13] and the derivation of rate laws [14, 15]. As model construction is challenging and as the information available in databases is limited, in general, a collection of candidate models can be constructed to compensate flaws in individual models [16]. For all these model candidates the parameters have to be estimated from experimental data, a challenging and usually ill-posed problem [17].

To determine maximum likelihood (ML) and maximum a posteriori (MAP) estimates for model parameters, high-dimensional nonlinear and non-convex optimization problems have to be solved.The non-convexity of the optimization problem poses challenges, such as local minima, which have to be addressed by the selection of optimization methods. Commonly used global optimization methods are multi-start local optimization [18], evolutionary and genetic algorithms [19], particle swarm optimizers [20], simulated annealing [21] and hybrid optimizers [22, 23] (see [24, 25, 26, 18] for a comprehensive survey). For ODE models with a few hundred parameters and state variables multi-start local optimization methods [18] and related hybrid methods [27] have proven to be successful. These optimization methods use the gradient of the objective function to establish fast local convergence. While the convergence of gradient based optimizers can be significantly improved by providing exact gradients (see e.g. [18, 28, 29]), the gradient calculation is often the computationally most demanding step.

The gradient of the objective function is usually approximated by finite differences. As this method is neither numerically robust nor computationally efficient, several parameter estimation toolboxes employ forward sensitivity analysis. This decreases the numerical error and computation time [18]. However, the dimension of the forward sensitivity equations increases linearly with both the number of state variables and parameters, rendering its application for genome-scale models problematic. In other research fields such as mathematics and engineering, adjoint sensitivity analysis is used for parameter estimation in ordinary and partial differential equation models. Adjoint sensitivity analysis is known to be superior to the forward sensitivity analysis when the number of parameters is large [30]. Adjoint sensitivity analysis has been used for inference of biochemical reaction networks [31, 32, 33]. However, the methods were never picked up by the systems and computational biology community, supposedly due to the theoretical complexity of adjoint methods, a missing evaluation on a set of benchmark models, and an absence of an easy-to-use toolbox.

In this manuscript, we provide an intuitive description of adjoint sensitivity analysis for parameter estimation in genome-scale biochemical reaction networks. We describe the end value problem for the adjoint state in the case of discrete-time measurement and provide an user-friendly implementation to compute it numerically. The method is evaluated on seven medium-to large-scale models. By using adjoint sensitivity analysis, the computation time for calculating the objective function gradient becomes effectively independent of the number of parameters with respect to which the gradient is evaluated. Furthermore, for large-scale models adjoint sensitivity analysis can be multiple orders of magnitude faster than other gradient calculation methods used in systems biology. The reduction of the time for gradient evaluation is reflected in the computation time of the optimization. This renders parameter estimation for large-scale models feasible on standard computers, as we illustrate for a comprehensive kinetic model of ErbB signaling.

## Methods

In this section we introduce the model class and the corresponding estimation problem. Subsequently, gradient calculation using finite differences, forward sensitivity analysis and adjoint sensitivity analysis is described and the theoretical complexity as well as some aspects of the numerical implementation are discussed.

### Mathematical model and experimental data

We consider ODE models for biochemical reaction networks,

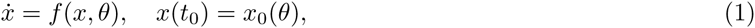
in which *x*(*t, θ*) ∈ ℝ^n_x_^ is the concentration vector at time *t* and *θ* ∈ ℝ^n_θ_^ denotes the parameter vector. Parameters are usually kinetic constants, such as binding affinities as well as synthesis, degradation and dimerization rates. The vector field ƒ: ℝ^n_x_^ × ℝ^n_θ_^ ↦ ℝ^n_x_^ describes the temporal evolution of the concentration of the biochemical species. The mapping *x*_*0*_: ℝ^n_θ_^ ↦ ℝ^n_x_^ provides the parameter dependent initial condition at time *t*_*0*_.

As available experimental techniques usually do not provide measurements of the concentration of all biochemical species, we consider the output map *h*: ℝ^n_x_^ × ℝ^n_θ_^ ↦ ℝ^n_y_^. This map models the measurement process, i.e. the dependence of the output (or observables) *y*(*t, θ*)∈ ℝ^n_y_^ at time point *t* on the state variables and the parameters,

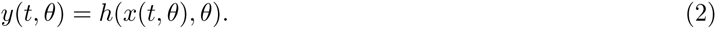

The *i*-th observable *yi* can be the concentration of a particular biochemical species (e.g. *y*_*i*_ = *x*_*l*_) as well as a function of several concentrations and parameters (e.g. *y*_*i*_ = θ_*m*_(*xl*_*1*_ + *xl*_*2*_)).

We consider discrete-time, noise corrupted measurements

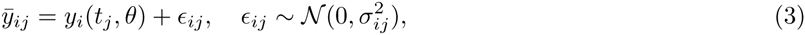
yielding the experimental data 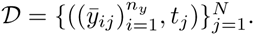 The number of time points at which measurements have been collected is denoted by *N.*

**Remark:** For simplicity of notation we assume throughout the manuscript that the noise variances, 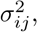, are known and that there are no missing values. However, the methods we will present in the following as well as the respective implementations also work if this is not the case. For details we refer to the *Supporting Information S1.*

### Maximum likelihood (ML) estimation

We estimate the unknown parameter *θ* from the experimental data 𝒟 using ML estimation. Parameters are estimated by minimizing the negative log-likelihood, an objective function indicating the difference between experiment and simulation. In the case of independent, normally distributed measurement noise with known variances the objective function is given by

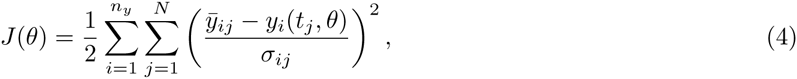
where *yi*(*t*_*j*_, *θ*) is the value of the output computed from (1) and (2) for parameter value *θ*. The minimization,

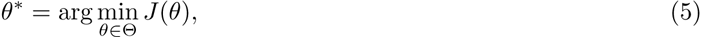
of this weighted least squares *J* yields the ML estimate of the parameters.

The optimization problem (5) is in general nonlinear and non-convex. Thus, the objective function can possess multiple local minima and global optimization strategies need to be used. For ODE models multi-start local optimization has been shown to perform well [18]. In multi-start local optimization, independent local optimization runs are initialized at randomly sampled initial points in parameter space. The individual local optimizations are run until the stopping criteria are met and the results are collected. The collected results are visualized by sorting them according to the final objective function value. This visualization reveals local optima and the size of their basin of attraction. For details we refer to the survey by Raue *et al.* [18]. In this study, initial points are generated using latin hypercube sampling and local optimization is performed using the interior point and the trust-region-reflective algorithm implemented in the MATLAB function fmincon.m. Gradients are computed using finite differences, forward sensitivity analysis or adjoint sensitivity analysis.

### Finite differences

A näive approximation to the gradient of the objective function with respect to *θ k* is obtained by finite differences,

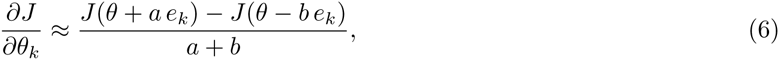
with *a, b* ≥ 0 and the kth unit vector *e*_*k*_. In practice forward differences (*a* = ∊, *b* = 0), backward differences (*a* = 0, *b* = ∊) and central differences (*a* = ∊, *b* = ∊) are widely used. For the computation of forward finite differences, this yields a procedure with three steps:

**Step 1** The state trajectory *x*(*t, θ*) and output trajectory *y*(*t, θ*) are computed.

**Step 2** The state trajectories *x*(*t, θ*(^*k*^)) and the output trajectories *y*(*t, θ*(^*k*^)) are computed for the perturbed parameters *θ*^k^ = *θ* + ∊e_*k*_ for *k* = 1,…, *n*_*θ.*_

**Step 3** The objective function gradient elements 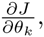 are computed from the output trajectory *y*(*t, θ*) and the perturbed output trajectory *y*(*t, θ*(^*k*^)) for *k* = 1,…, *n*_*θ.*_

In theory, forward and backward differences provide approximations of order e while central differences provide more accurate approximations of order *∊*_*2*_, provided that *J* is sufficiently smooth. In practice the optimal choice of *a* and *b* depends on the accuracy of the numerical integration [18]. If the integration accuracy is high, an accurate approximation of the gradient can be achieved using *a, b* ≪ 1. For lower integration accuracies, larger values of *a* and *b* usually yield better approximations. A good choice of *a* and *b* is typically not clear *a priori* (cf. [34] and the references therein).

The computational complexity of evaluating gradients using finite differences is affine linear in the number of parameters. Forward and backward differences require in total *n*_*θ*_ + 1 function evaluations. Central differences require in total *2 n*_*θ*_ function evaluations. As already a single simulation of a large-scale model is time-consuming, the gradient calculation using finite differences can be limiting.

### Forward sensitivity analysis

State-of-the-art systems biology toolboxes, such as the MATLAB toolbox Data2Dynamics [7], use forward sensitivity analysis for gradient evaluation. The gradient of the objective function is

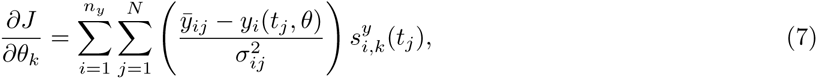
with 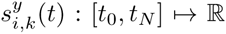 denoting the sensitivity of output *y*_*i*_ at time point *t* with respect to parameter θ_k_. Governing equations for the sensitivities are obtained by differentiating (1) and (2) with respect to *θk* and reordering the derivatives. This yields

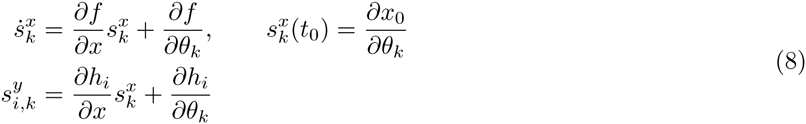
with 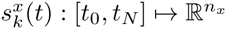 denoting the sensitivity of the state *x* with respect to *θk.* Note that here and in the following, the dependencies of ƒ, *h*, *x*_*0*_ and their (partial) derivatives on *t, x* and *θ* are not stated explicitly but have the to be assumed. For a more detailed presentation we refer to the *S1 Supporting Information Section 1.*

Forward sensitivity analysis consists of three steps:

**Step 1** The state trajectory *x*(*t, θ*) and output trajectory *y*(*t, θ*) are computed.

**Step 2** The state sensitivities 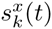 and the output sensitivities 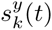 are computed using the state trajectory *x*(*t, θ*) for *k* = 1,…, *n*_*θ*_.

**Step 3** The objective function gradient elements 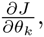 are computed from the output sensitivities 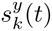 and the output trajectory *y*(*t, θ*) for *k* = 1,…, *n*_*θ*_

Step 1 and 2 are often combined, which enables simultaneous error control and the reuse of the Jacobian [30]. The simultaneous error control allows for the calculation of accurate and reliable gradients. The reuse of the Jacobian improves the computational efficiency.

The number of state and output sensitivities increases linearly with the number of parameters. While this is unproblematic for small-and medium-sized models, solving forward sensitivity equations for systems with several thousand state variable bears technical challenges. Code compilation can take multiple hours and require more memory than what is available on standard machines. Furthermore, while forward sensitivity analysis is usually faster than finite differences, in practice the complexity still increases roughly linearly with the number of parameters.

### Adjoint sensitivity analysis

In the numerics community, adjoint sensitivity analysis is frequently used to compute the gradients of a functional with respect to the parameters if the function depends on the solution of a differential equation [35]. In contrast to forward sensitivity analysis, adjoint sensitivity analysis does not rely on the state sensitivities 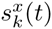 but on the adjoint state *p*(*t*).

The calculation of the objective function gradient using adjoint sensitivity analysis consists of three steps:

**Step 1** The state trajectory *x*(*t, θ*) and output trajectory *y*(*t, θ*) are computed.

**Step 2** The trajectory of the adjoint state *p*(*t*) is computed.

**Step 3** The objective function gradient elements 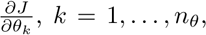 are computed from the state trajectory *x*(*t, θ*), the adjoint state trajectory *p*(*t*) and the output trajectory *y*(*t, θ*).

Step 1 and 2, which are usually the computationally intensive steps, are independent of the parameter dimension. The complexity of Step 3 increases linearly with the number of parameters, yet the computation time required for this step is typically negligible.

The calculation of state and output trajectories (Step 1) is standard and does not require special methods. The non-trivial element in adjoint sensitivity analysis is the calculation of the adjoint state *p*(*t*) ∈ ℝ^n_x_^ (Step 2). For discrete-time measurements - the usual case in systems and computational biology - the adjoint state is piece-wise continuous in time and defined by a sequence of backward differential equations. For *t* > t_N_, the adjoint state is zero, *p*(*t*) = 0. Starting from this end value the trajectory of the adjoint state is calculated backwards in time, from the last measurement *t* = *t*_*N*_ to the initial time *t* = *t*_*0*_. At the time points at which measurements have been collected, *t*_*N*_,…,*t*_1_, the adjoint state is reinitialised as

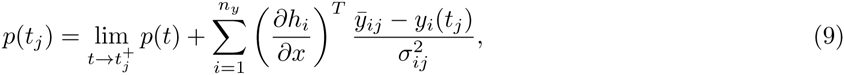
which usually results in a discontinuity of *p*(*t*) at *t*_*j*_. Starting from the end value *p*(*t*_*j*_) as defined in (9) the adjoint state evolves backwards in time until the next measurement point *t*_*j—1*_ or the initial time to is reached. This evolution is governed by the time-dependent linear ODE

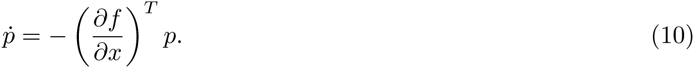

The repeated evaluation of (9) and (10) until *t* = *t*_*0*_ yields the trajectory of the adjoint state. Given this trajectory, the gradient of the objective function with respect to the individual parameters is

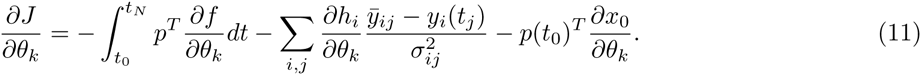

Accordingly, the availability of the adjoint state simplifies the calculation of the objective function to *nθ* one-dimensional integration problems over short time intervals whose union is the total time interval [*t*_*0*_,*t*_*N*_].

#### Algorithm 1: Gradient evaluation using adjoint sensitivity analysis

*% State and output*

*Step 1* Compute state and output trajectories using (1) and (2).

*% Adjoint state*

*Step 2.1Set* end value for adjoint state, ∀*t* > *p*(*t*) = 0.

**for** *j* = *N to 1* **do**

|*Step 2.2*Compute end value for adjoint state according to the *j*th measurement using (9).

|*Step 2.3*Compute trajectory of adjoint state on time interval *t* = (*t*_*j-1*_,*t*_*j*_] by solving (10).

**end**

*% Objective function gradient*

**for** *k =* 1 *to n*_*θ*_ **do**

|*Step 3* Evaluation of the sensitivity *∂J*/*∂θ*_*k*_ using (11). **end**

Pseudo-code for the calculation of the adjoint state and the objective function gradient is provided in Algorithm 1. We note that in order to use standard ODE solvers the end value problem (10) can be transformed in an initial value problem by applying the time transformation *τ* = *t*_*N*_ — *t.* The derivation of the adjoint sensitivities for discrete-time measurements is provided in the *S1 Supporting Information Section 1.*

The key difference of the adjoint compared to the forward sensitivity analysis is that the derivatives of the state and the output trajectory with respect to the parameters are not explicitly calculated. Instead, the sensitivity of the objective function is directly computed. This results in practice in a computation time of the gradient which is almost independent of the number of parameters. A visual summary of the different sensitivity analysis methods is provided in Figure 1. Besides the procedures also the computational complexity is indicated.

**Figure 1:**
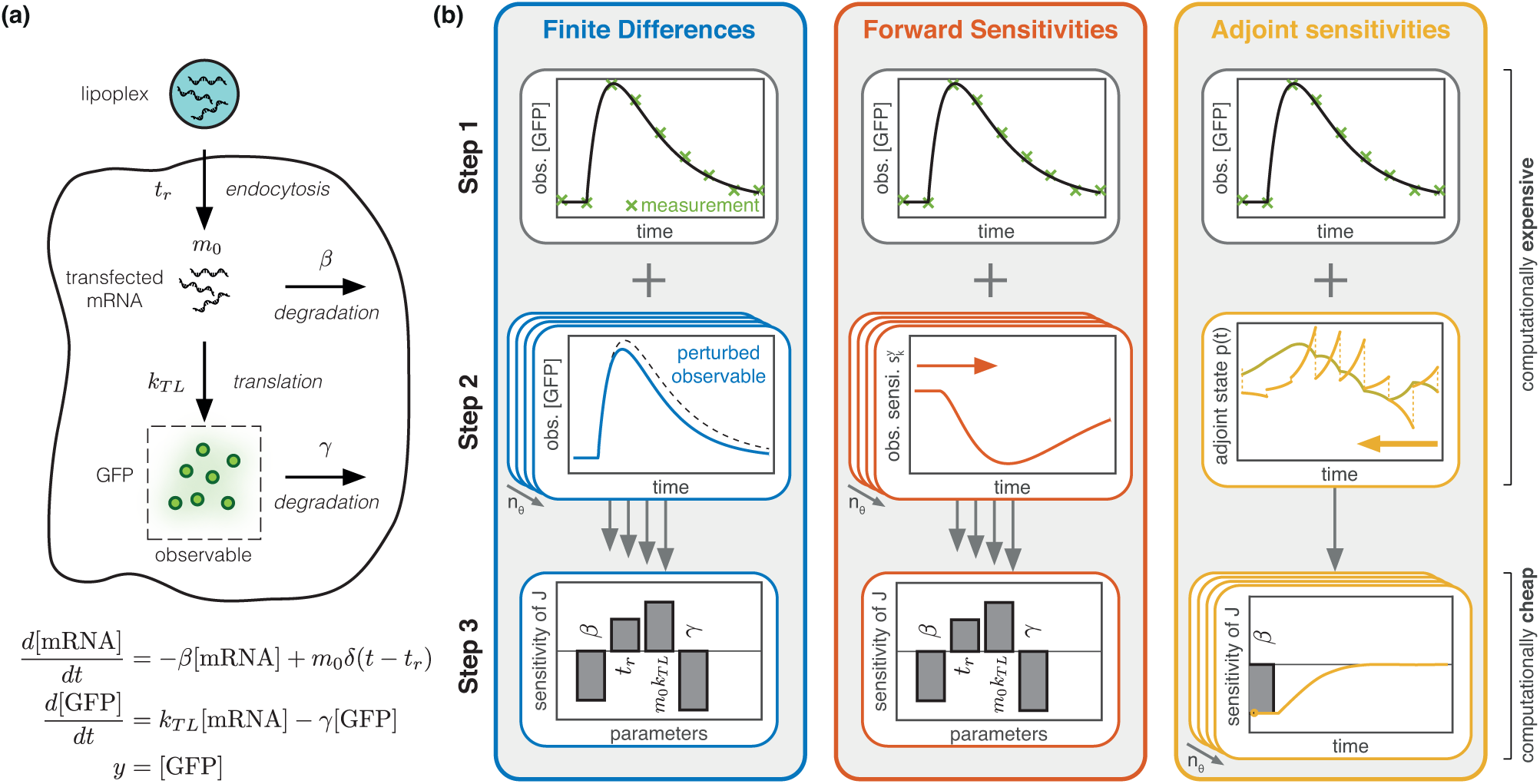
Illustration of gradient calculation using finite differences, forward sensitivity analysis and adjoint sensitivity equations for a model of mRNA transfection. (a) Sketch and mathematical formulation of the mathematical model of mRNA transfection presented by [36]. The intracellular release of mRNA at time point *t*_*r*_ is modeled using the Dirac delta distribution *δ.* (b) Illustration of finite differences, forward sensitivity analysis and adjoint sensitivity analysis for the model of mRNA transfection: (top) Step 1: simulation of model; (middle) Step 2: intermediate step for gradient calculation; and (bottom) Step 3: calculation of gradient from intermediate results. For all methods, Step 1 and 2 involve numerical simulation (the direction indicated by the arrow) and are computationally demanding, while Step 3 is computationally negligible.

### Implementation

The implementation of adjoint sensitivity analysis is non-trivial and error-prone. To render this method available to the systems and computational biology community, we implemented the Advanced Matlab Interface for CVODES and IDAS (AMICI). This toolbox allows for a simple symbolic definition of ODE models (1)-(2) as well as the automatic generation of native C code for efficient numerical simulation. The compiled binaries can be executed from MATLAB for the numerical evaluation of the model and the objective function gradient. Internally, the SUNDIALS solvers suite is employed [30], which offers a broad spectrum of state-of-the-art numerical integration of differential equations. In addition to the standard functionality of SUNDIALS, our implementation allows for parameter and state dependent discontinuities. The toolbox and a detailed documentation can be downloaded from ICB-DCM.github.io/AMICI/.

## Results

In the following, we will illustrate the properties of adjoint sensitivity analysis for biochemical reaction networks. For this purpose, we study several models provided in the BioPreDyn benchmark suite [27] and from the curated branch of the Biomodels Database [37]. We compare adjoint sensitivity analysis with forward sensitivity analysis and finite differences regarding accuracy, computational efficiency and scalability for a set of medium-to large-scale models.

### Investigated Models

For the comparison of different gradient calculation methods, we consider a set of standard models from the Biomodels Database [37] and the BioPreDyn benchmark suite [27]. From the biomodels database we considered models for the regulation of insulin signaling by oxidative stress (BM1) [38], the sea urchin endomesoderm network (BM2) [39], and the ErbB sigaling pathway (BM3) [40]. From BioPreDyn benchmark suite we considered models for central carbon metabolism in E. coli (B2) [41], enzymatic and transcriptional regulation of carbon metabolism in E. coli (B3) [42], metabolism of CHO cells (B4) [43], and signaling downstream of EGF and TNF (B5) [44]. Genome-wide kinetic metabolic models of S. cerevisiae and E.coli (B1) [45] contained in the BioPreDyn benchmark suite and the Biomodels Database [15, 45] were disregarded due to previously reported numerical problems [45, 27]. The considered models possess 18-500 state variable and 86-1801 parameters. A comprehensive summary regarding the investigated models is provided in Table 1.

**Table 1:**
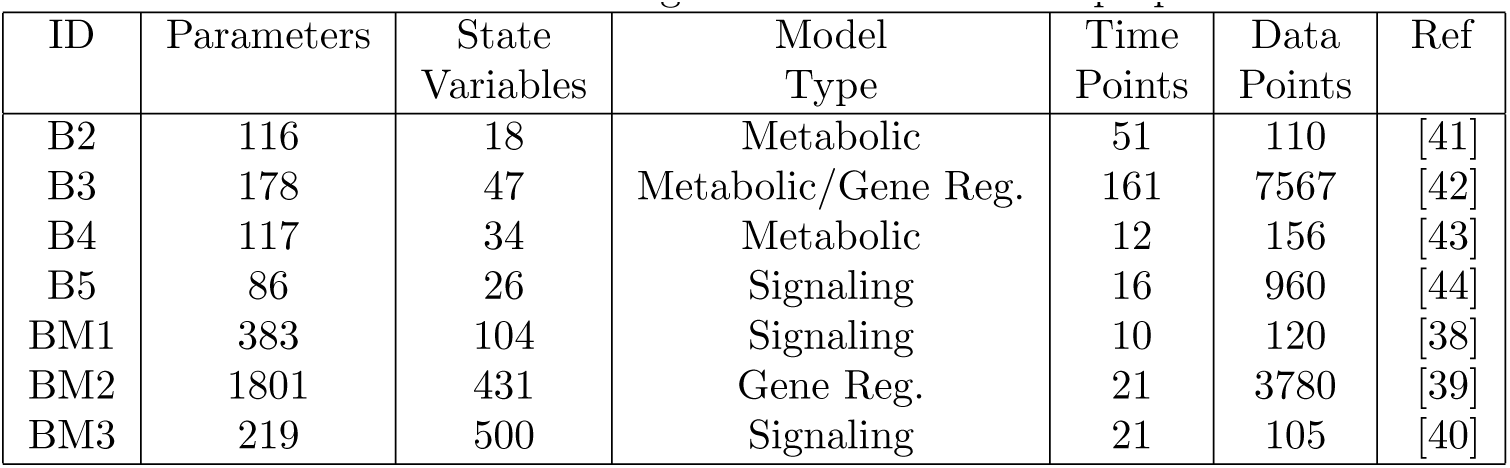
List of investigated models and their properties

| ID | Parameters | State Variables | Model Type | Time Points | Data Points | Ref |
| --- | --- | --- | --- | --- | --- | --- |
| B2 | 116 | 18 | Metabolic | 51 | 110 | [41] |
| B3 | 178 | 47 | Metabolic/Gene Reg. | 161 | 7567 | [42] |
| B4 | 117 | 34 | Metabolic | 12 | 156 | [43] |
| B5 | 86 | 26 | Signaling | 16 | 960 | [44] |
| BM1 | 383 | 104 | Signaling | 10 | 120 | [38] |
| BM2 | 1801 | 431 | Gene Reg. | 21 | 3780 | [39] |
| BM3 | 219 | 500 | Signaling | 21 | 105 | [40] |

To obtain realistic simulation times for adjoint sensitivities realistic experimental data is necessary (see S1Supporting Information Section 3). For the BioPreDyn models we used the data provided in the suite, for the ErbB signaling pathway we used the experimental data provided in the original publication and for the remaining models we generated synthetic data using the nominal parameter provided in the SBML definition.

In the following, we will compare the performance of forward and adjoint sensitivities for these models. As the model of ErbB signaling has the largest number of state variables and is of high practical interest in the context of cancer research, we will analyze the scalability of finite differences and forward and adjoint sensitivity analysis for this model in greater detail. Moreover, we will compare the computational efficiency of forward and adjoint sensitivity analysis for parameter estimation for the model of ErbB signaling.

### Scalability of gradient evaluation using adjoint sensitivity analysis

The evaluation of the objective function gradient is the computationally demanding step in deterministic local optimization. For this reason, we compared the computation time for finite differences, forward sensitivity analysis and adjoint sensitivity analysis and studied the scalability of these approaches at the nominal parameter *θ*_0_ which was provided in the SBML definitions of the investigated models.

For the comprehensive model of ErbB signaling we found that the computation times for finite differences and forward sensitivity analysis behave similarly (Figure 2a). As predicted by the theory for both methods the computation time increased linearly with the number of parameters. Still, forward sensitivities are computationally more efficient than finite differences, as reported in previous studies [18].

**Figure 2:**
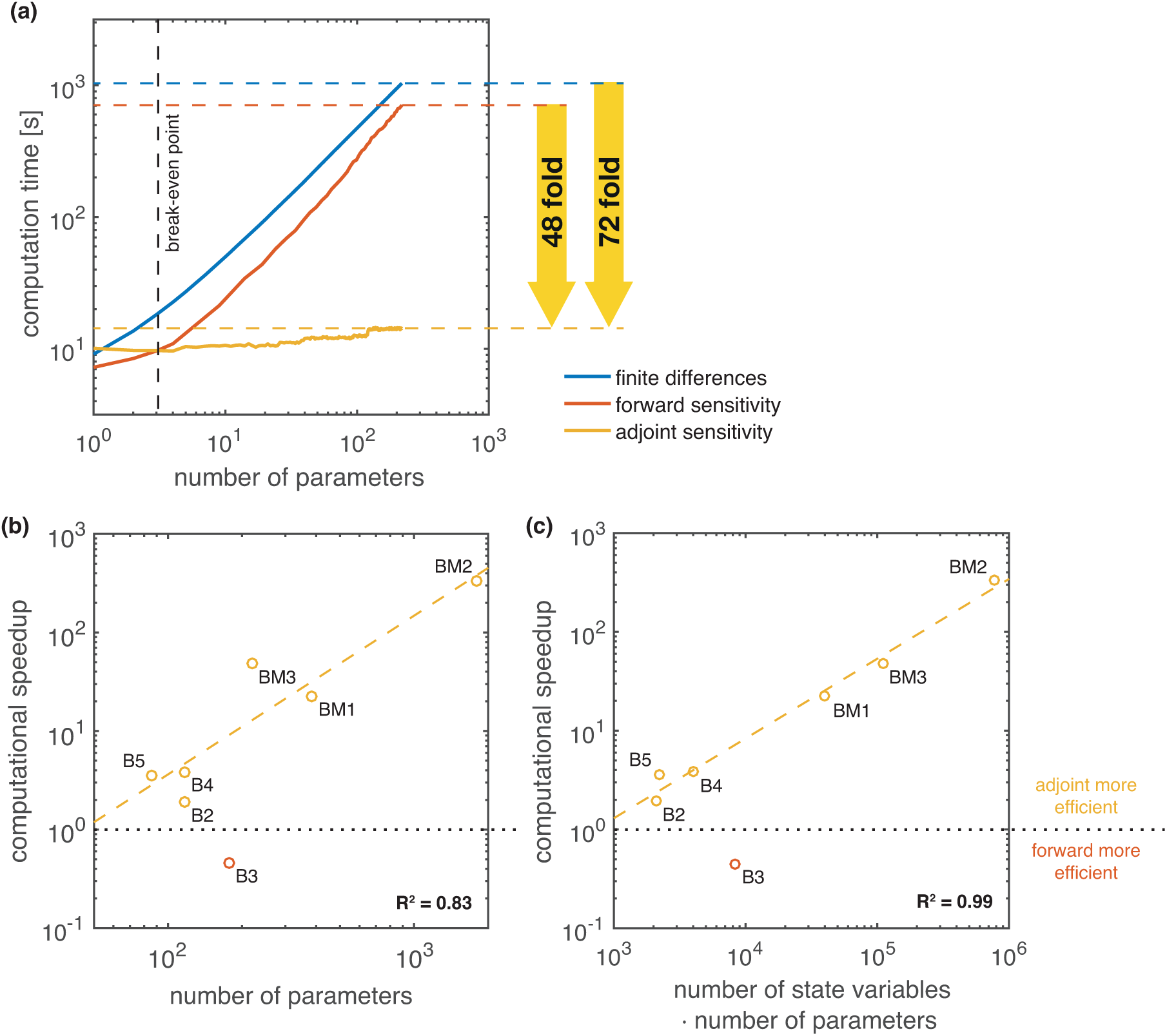
Comparison of gradient computation times for finite differences and forward and adjoint sensitivity analysis. (a) Scaling of computation time with respect to the number of parameters for the model of ErbB signaling (BM3). Computation time for finite differences and forward sensitivity equations increases roughly linearly. Computation time for adjoint sensitivity analysis is almost independent of the number of parameters but possesses a higher initial cost. Adjoint sensitivity analysis is 48 times faster than forward sensitivity analysis when considering all parameters. (b,c) Speedup when using adjoint sensitivity analysis over forward sensitivity analysis for gradient computation evaluated for all investigated models compared against *n*_*θ*_ and *n*_*x*_. *n*_*θ*_. Regression curves (dashed lines) have been fitted to the results of all models excluding B3, which seems to be an outlier. All computations were performed on a MacBook Pro with an 2.9 GHz Intel Core i7 processor.

Adjoint sensitivity analysis requires the solution to the adjoint problem, independent of the number of parameters. For the considered model, solving the adjoint problem a single time takes roughly 2-3-times longer than solving the forward problem. Accordingly, adjoint sensitivity analysis with respect to a small number of parameter is disadvantageous. However, adjoint sensitivity analysis scales better than forward sensitivity analysis and finite differences. Indeed, the computation time for adjoint sensitivity analysis is almost independent of the number of parameters. While computing the sensitivity with respect to a single parameter takes on average 10.09 seconds, computing the sensitivity with respect to all 219 parameters takes merely 14.32 seconds. We observe an average increase of 1.9 · 10-^2^ seconds per additional parameter for adjoint sensitivity analysis which is significantly lower than the expected 3.24 seconds for forward sensitivity analysis and 4.72 seconds for finite differences. If the sensitivities with respect to more than 4 parameters are required, adjoint sensitivity analysis outperforms both forward sensitivity analysis and finite differences. For 219 parameters, adjoint sensitivity analysis is 48-times faster than forward sensitivities and 72-times faster than finite differences.

To ensure that the observed speedup is not unique to the model of ErbB signaling (BM3) we also evaluated the speedup of adjoint sensitivity analysis over forward sensitivity analysis on models B2-5 and BM1-2. The results are presented in Figure 2b,c. We find that for all models, but model B3, gradient calculation using adjoint sensitivity is computationally more efficient than gradient calculation using forward sensitivities (speedup ¿ 1). For model B3 the backwards integration required a much higher number of integration steps (4 · 10^6^) than the forward integration (6 · 10^3^), which results to a poor performance of the adjoint method. One reason for this poor performance could be that, in contrast to other models, the right hand side of the differential equation of model B3 consists almost exclusively of non-linear, non-mass-action terms.

Excluding model B3 we find an polynomial increase in the speedup with respect to the number of parameters *n*_*θ*_ (Figure 2b), as predicted by theory. Moreover, we find that the product *n*_*θ*_ · *n*_*x*_ which corresponds to the size of the system of forward sensitivity equations, is an even better predictor (*R*^2^ = 0.99) than *n θ* alone (*R*^2^ = 0.83). This suggest that adjoint sensitivity analysis is not only beneficial for systems with a large number of parameters, but can also be beneficial for systems with a large number of state variables. As we are not aware of any similar observations in the mathematics or engineering community, this could be due to the structure of biological reaction networks.

Our results suggest that adjoint sensitivity analysis is an excellent candidate for parameter estimation in large-scale models as it provides good scaling with respect to both, the number of parameters and the number of state variables.

### Accuracy and robustness of gradients computing adjoint sensitivity analysis

Efficient local optimization requires accurate and robust gradient evaluation [18]. To assess the accuracy of the gradient computed using adjoint sensitivity analysis, we compared this gradient to the gradients computed via finite differences and forward sensitivity analysis. Figure 3 visualizes the results for the model of ErbB signaling (BM3) at the nominal parameter θ_0_ which was provided in the SBML definition. The results are similar for other starting points.

**Figure 3:**
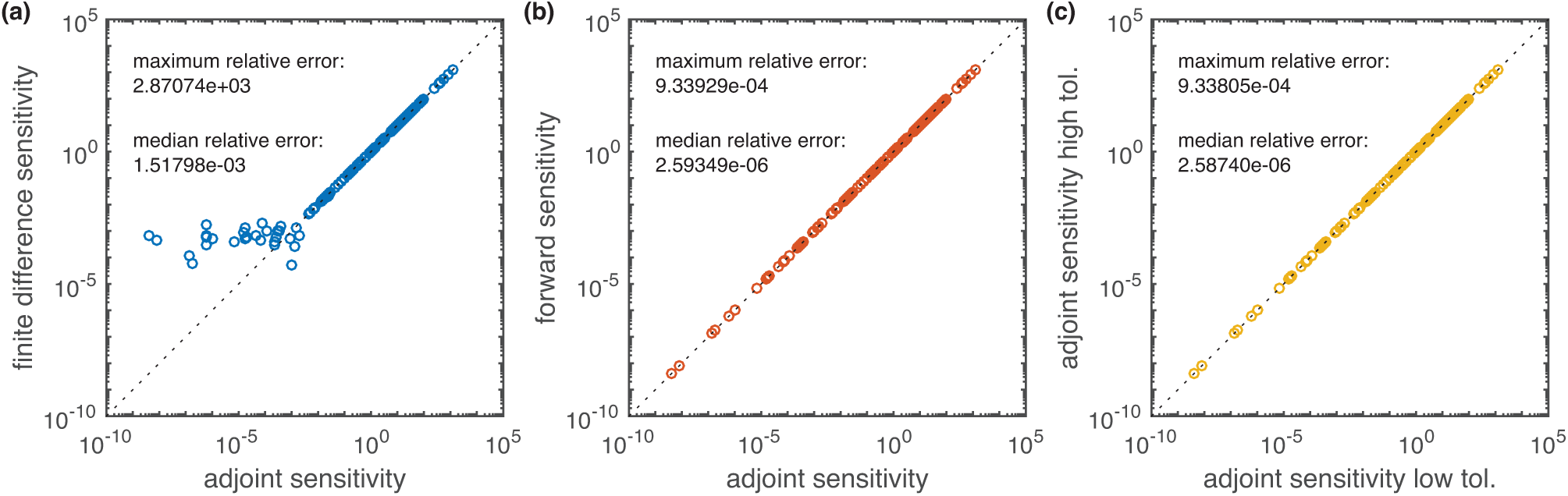
Comparison of the gradients computed using adjoint sensitivity equations with gradients computed using finite differences and forward sensitivity equations with default accuracies (absolute error < 10-^16^, relative error < 10-^8^). Each point represents the absolute value of one gradient element. Points on the diagonal indicate a good agreement. (a) Forward finite differences with ∊ = 10-^3^ vs. adjoint sensitivities. (b) Forward sensitivities vs. adjoint sensitivities. (c) Adjoint sensitivities with high accuracies (absolute error < 10-^32^, relative error < 10-^16^) and default accuracies (absolute error < 10-^16^, relative error < 10-^8^).

The comparison of the gradients obtained using finite differences and adjoint sensitivity analysis revealed small discrepancies (Figure 3a). The median relative difference (as defined in S1 Supporting Information Section 2) between finite differences and adjoint sensitivity analysis is 1.5 · 10-^3^. For parameters θ_k_ to which the objective function *J* was relatively insensitive, *δJ*/*δθ*_*k*_ < 10-^2^, there are much higher discrepancies, up to a relative error of 2.9 · 10^3^.

Forward and adjoint sensitivity analysis yielded almost identical gradient elements over several orders of magnitude (Figure 3b). This was expected as both forward and adjoint sensitivity analysis exploit error-controlled numerical integration for the sensitivities. To assess numerical robustness of adjoint sensitivity analysis, we also compared the results obtained for high and low integration accuracies (Figure 3c). For both comparisons we found the similar median relativer and maximum relative error, namely 2.6·10^−6^ and 9.3·10^−4^. This underlines the robustness of the sensitivitity based methods and ensures that differences observed in Figure 3a indeed originate from the inaccuracy of finite differences.

Our results demonstrate that adjoint sensitivity analysis provides objective function gradients which are as accurate and robust as those obtained using forward sensitivity analysis.

### Optimization of large-scale models using adjoint sensitivity analysis

As adjoint sensitivity analysis provides accurate gradients for a significantly reduced computational cost, this can boost the performance of a variety of optimization methods. Yet, in contrast to forward sensitivity analysis, adjoint sensitivities do not yield sensitivities of observables and it is thus not possible to approximate the Hessian of the objective function via the Fisher Information Matrix [46]. This prohibits the use of possibly more efficient Newton-type algorithms which exploit second order information. Therefore, adjoint sensitivities are limited to quasi-Newton type optimization algorithms, e.g. the Broyden-Fletcher-Goldfarb-Shanno (BFGS) algorithm [47, 48], for which the Hessian is iteratively approximated from the gradient during optimization. In principle, the exact calculation of the Hessian and Hessian-Vector products is possible via second order forward and adjoint sensitivity analysis [49, 50], which possess similar scaling properties as the first order methods. However, both forward and adjoint approaches come at an additional cost and are thus not considered in this study.

To assess whether the use of adjoint sensitivities for optimization is still viable, we compared the performance of the interior point algorithm using adjoint sensitivity analysis with the BFGS approximation of the Hessian to the performance of the trust-region reflective algorithm using forward sensitivity analysis with Fisher Information Matrix as approximation of the Hessian. For both algorithms we used the MATLAB implementation in fmincon.m. The employed setup of the trust-region algorithm is equivalent to the use of lsqnonlin.m which is the default optimization algorithm in the MATLAB toolbox Data2Dynamics [7], which was employed to win several DREAM challenges. For the considered model the computation time of forward sensitivities is comparable in Data2Dynamics and AMICI. Therefore, we expect that Data2Dynamics would perform similar to the trust-region reflective algorithm coupled to forward sensitivity analysis.

We evaluated the performance for the model of ErbB signaling based on 100 multi-starts which were initialized at the same initial points for both optimization methods. For 41 out of 100 initial points the gradient could not be evaluated due numerical problems. These optimization runs are omitted in all further analysis. To limit the expected computation to a bearable amount we allowed a maximum of 10 iterations for the forward sensitivity approach and 500 iterations for the adjoint sensitivity approach. As the previously observed speedup in gradient computation was roughly 48 fold, we expected this setup should yield similar computation times for both approaches.

We found that for the considered number of iterations, both approaches perform similar in terms of objective function value compared across iterations (Figure 4a,b). However, the computational cost of one iteration was much cheaper for the optimizer using adjoint sensitivity analysis. Accordingly, given a fixed computation time the interior-point method using adjoint sensitivities outperforms the trust-region method employing forward sensitivities and the FIM (Figure 4c,d). In the allowed computation time, the interior point algorithm using adjoint sensitivities could reduce the objective function by up to two orders of magnitude (Figure 4c). This was possible although many model parameters seem to be non-identifiable (see S1 Supplemtary Information Section 4), which can cause problems.

**Figure 4:**
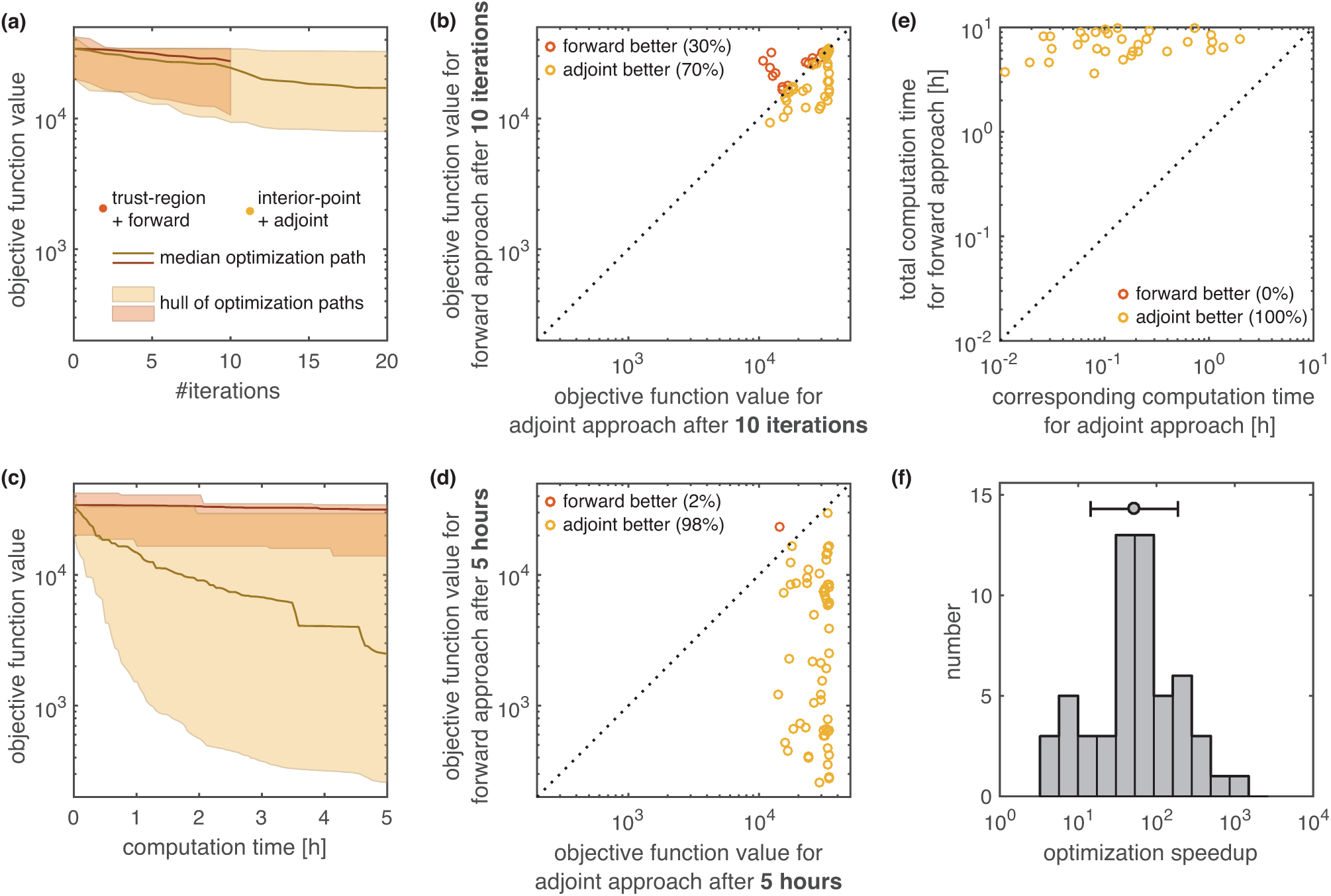
Comparison of optimization speed using forward and adjoint sensitivities for the model of ErbB signaling. For local optimization using forward sensitivity analysis (trust-region method) and local optimization using adjoint sensitivity analysis (interior-point method) we quantified the computation time across 100 local optimization runs with different initial conditions. For 41 out of 100 initial points the gradient could not be evaluated due to numerical problems. These optimization runs are omitted in all further analysis. (a,c) Comparison of objective function value with respect to iteration number and computation time. The hulls and medians computed for both methods are depicted as shaded areas and solid lines. (b,d) Pairwise comparison of objective function value after 10 iterations and 5 hours for both methods. Each dot corresponds to one initial point for the optimization. The coloring indicates which method performed better. (e) Pairwise comparison of the time required to reach the final objective function value achieved in the forward approach. For the adjoint approach the equivalent time is the minimal time to reach the same objective function value. Each dot corresponds to one initial point for the optimization. (f) Histogram of speedup by using adjoint sensitivity analysis over forward sensitivity analysis for individual initial points, computed from (e). All computations were performed on a linux cluster. Runs with same initial conditions were carried out on the same computation node.

To quantify the speedup of the optimization using adjoint sensitivity analysis over the optimization using forward sensitivity analysis, we performed a pairwise comparison of the minimal time required by the adjoint sensitivity approach to reach the final objective function value of the forward sensitivity approach for the individual points (Figure 4e). The median speedup achieved across all multi-starts was 54 (Figure 4f), which was similar to the 48 fold speedup achieved in the gradient computation. The availability of the Fisher Information Matrix for forward sensitivities did not compensate for the significantly reduced computation time achieved using adjoint sensitivity analysis. This could be due to the fact that adjoint sensitivity based approach, being able to carry out many iterations in a short time-frame, can build a reasonable approximation of the Hessian approximation relatively fast.

In summary, this application demonstrates the applicability of adjoint sensitivity analysis for parameter estimation in large-scale biochemical reaction networks. Possessing similar accuracy as forward sensitivities, the scalability is improved which results in an increased optimizer efficiency. For the model of ErbB signaling, optimization using adjoint sensitivity analysis outperformed optimization using forward sensitivity analysis.

## Discussion

Mechanistic mathematical modeling at the genome scale is an important step towards a holistic understanding of biological processes. To enable modeling at this scale, scalable computational methods are required which are applicable to networks with thousands of compounds. In this manuscript, we present a gradient computation method which meets this requirement and which renders parameter estimation for large-scale models significantly more efficient. Adjoint sensitivity analysis, which is extensively used in other research fields, is a powerful tool for estimating parameters of large-scale ODE models of biochemical reaction networks.

Our study of several benchmark models with up to 500 state variables and up to 1801 parameters demonstrated that adjoint sensitivity analysis provides accurate gradients in a computation time which is much lower than for established methods and effectively independent of the number of parameters. To achieve this, the adjoint state is computed using a piece-wise continuous backward differential equation. This backward differential equation has the same dimension as the original model, yet the computation time required to solve it usually is slightly larger. As a result, finite differences and forward sensitivity analysis might be more efficient if the sensitivities with respect to a few parameters are required. The same holds for alternatives like complex-step derivative approximation techniques [51] and forward-mode automatic differentiation [52, 28]. For systems with many parameters, adjoint sensitivity analysis is advantageous. A scalable alternative might be reverse-mode automatic differentiation [53, 28], which remains to be evaluated for the considered class of problems.

For the model of ErbB signaling we could show that adjoint sensitivity based optimization outperforms forward sensitivity based optimization, which is the standard in most systems biology toolboxes. With the availability of the MATLAB toolbox AMICI the adjoint sensitivity based approach becomes accessible for other researchers. AMICI allows for the fully automated generation of executables for adjoint or forward sensitivity analysis from symbolic model definitions. This way, the toolbox is easy-to-use and can easily be integrated with existing toolboxes. Also other MATLAB toolboxes for computational modeling, e.g. AMIGO [6], Data2Dynamics [7], MEIGO [54] and SBtoolbox2 [55] could be extended to exploit adjoint sensitivity analysis. In addition to adjoint sensitivity analysis, these MATLAB toolboxes could exploit forward sensitivity analysis available via AMICI, as AMICI yields computation times comparable to those of tailored numerical methods such as odeSD [56] (S1 Supporting Information Section 5) or Data2Dynamics [7]. Moreover AMICI comes with detailed documentation and is already now used by several research labs.

Our study of the model of ErbB signaling suggests that for the available data, a large number of parameters remains non-identifiable. While novel technologies provide rich dataset, we expect that non-identifiably will remain a problem. In particular if merely relative measurements are available, as the case for many measurement techniques, additional unknown scaling factors need to be introduced. These scaling factors are, in combination with initial conditions and total abundances, often the source of practical and structural non-identifiabilites [18]. Fortunately, for a broad range of biological questions, these information are not necessary and also state-of-the-art methods optimization seem to work reasonably well in the presence of non-identifiabilities. For the considered model of EreB signaling, we were able to achieve a significant decrease in the objective function value, despite the non-identifiability of parameters. This demonstrates that gradient based optimization is still feasible for large-scale problems. Yet, we believe that convergence of the optimizer could be improved by regularizing the objective function by integrating prior knowledge, possibly in a Bayesian framework [57], from databases such as SABIO-RK [58] or BRENDA [59].

Beyond the use in optimization, gradients computed using adjoint sensitivity analysis will also facilitate the development of more efficient uncertainty analysis methods. Riemann manifold Langevin and Hamiltonian Monte Carlo methods [60, 61] exploit the first and second order local structure of the posterior distribution and profit from more efficient gradient evaluation. The same holds for novel emulator-based sampling procedures [62] and approaches for posterior approximation [63]. By exploiting the proposed approach, rigorous Bayesian parameter estimation for models with hundreds of parameters could become a standard tool instead of an exception [64, 65].

In conclusion, adjoint sensitivity analysis will facilitate the development of large-and genome-scale mechanistic models for cellular processes as well as other (multi-scale) biological processes [66]. This will complement available statistical analysis methods for omics data [67] by providing mechanistic insights and render a holistic understanding feasible.

